# Identification of antibody-drug conjugate payloads which are substrates of ATP-binding cassette drug efflux transporters

**DOI:** 10.1101/2025.05.22.651305

**Authors:** Jacob S. Roth, Hui Guo, Lu Chen, Min Shen, Omotola Gbadegesin, Robert W. Robey, Michael M. Gottesman, Matthew D. Hall

## Abstract

**Aim:** Antibody-drug conjugates (ADCs) feature an antibody recognizing a specific protein joined to a potent toxic payload. Numerous antibody-drug conjugates have received FDA approval; however, clinical resistance arises. Resistance mechanisms include decreased expression or mutation of the antibody target, impaired payload release, or increased expression of ATP-binding cassette (ABC) efflux transporters associated with multidrug resistance. We therefore sought to characterize the interactions of ABC multidrug transporters with ADC payloads.

**Methods:** We performed a high-throughput screen with 27 common ADC payloads using cells lines expressing ABC transporters P-glycoprotein (P-gp, encoded by *ABCB1*) or ABCG2 (encoded by *ABCG2*). Confirmatory assays were also performed using cells transfected to express P-gp, ABCG2, or MRP1 (encoded by *ABCC1*).

**Results:** Several commonly used ADC payloads were substrates of P-gp, including calicheamicin gamma1, monomethyl auristatin E, DM1, and DM4. All the pyrrolobenzodiazepines tested—SJG136, SGD-1882, SG2057, and SG3199—were substrates of P-gp, ABCG2, and MRP1. The modified anthracyclines nemorubicin and its metabolite PNU-159682 were poorly transported by both ABCB1 and ABCG2 and displayed nanomolar to picomolar toxicity. Further, we found that the efficacy of the FDA-approved ADC mirvetuximab soravtansine, with DM4 as the toxic payload, was decreased in cell lines expressing P-gp. Duocarmycin DM and PNU-159682 were exquisitely toxic to a panel of 99 cancer cell lines of varying origins.

**Conclusion:** Several commonly used ADC payloads can be transported by ABC transporters, potentially leading to transporter-mediated drug resistance in patients. Future ADCs should be developed using payloads that are not ABC transporter substrates.

## Introduction

Antibody-drug conjugates (ADCs) have received significant attention as a modality for selectively targeting tumor cells. ADCs are composed of an antibody targeting a specific cell surface antigen conjugated to a small-molecule toxin (payload) via a cleavable linker^[1]^. Following systemic distribution and binding to the antigen, the ADC is internalized and processed within the target cell lysosomes, liberating the toxin to exert its cytotoxic effects. Thus, provided the antibody is targeted to a protein/epitope that is selectively expressed on the extracellular surface of cancer cells and minimally in other cell-types in the body, the ADC should selectively kill tumor cells^[2]^. The first ADC was granted FDA approval in 2000 for the treatment of acute myeloid leukemia (AML); gemtuzumab ozogamicin targets CD33 and carries N-acetyl-gamma-calicheamicin as the cytotoxic payload^[3]^. Since then, several other ADCs have been approved for the treatment of cancer including brentuximab vedotin for Hodgkin lymphoma^[4]^, trastuzumab emtansine for breast cancer^[5]^, trastuzumab deruxtecan for breast cancer^[6]^ (recently approved for any HER2-positive solid cancer^[7]^), loncastuximab tesirine for diffuse large B-cell lymphoma^[8]^, enfortumab vedotin for urothelial cancers^[9]^ and mirvetuximab soravtansine for cisplatin-resistant ovarian cancers^[10]^. FDA-approved ADCs, and ADCs in clinical trials, have employed highly cytotoxic natural product-derived payloads which cause cell death via various modalities including induction of DNA damage, disruption of microtubules, and inhibition of topoisomerase^[11]^. Many of these agents originate from molecules that were explored as small molecule (untargeted) chemotherapeutics, but potent cytotoxic side-effects led to their abandonment. A partial listing of approved ADCs and ADCs currently in clinical trials is included in Table 1.

**Table 1:** Approved and clinical candidate ADC small molecule payloads.

| Name | Brand Name | Payload | Linker | FDA Status |
| --- | --- | --- | --- | --- |
| Gemtuzumab ozogamicin | Mylotarg | Calicheamicin | Cleavable | Approved |
| Inotuzumab ozogamicin | Besponsa | Calicheamicin | Cleavable | Approved |
| Trastuzumab deruxtecan | Enhertu | Dxd (exatecan derivative) | Cleavable | Approved |
| Datopotamab deruxtecan | Datroway | Dxd (exatecan derivative) | Cleavable | Approved |
| Ado-trastuzumab emtansine | Kadcyla | DM1 (Maytansinoid analog) | Non-Cleavable | Approved |
| Mirvetuximab soravtansine | Elahere | DM4 (Maytansinoid analog) | Cleavable | Approved |
| Brentuximab vedotin | Adcetris | Monomethyl auristatin E (MMAE) | Cleavable | Approved |
| Polatuzumab vedotin | Polivy | Monomethyl auristatin E (MMAE) | Cleavable | Approved |
| Enfortumab vedotin | Padcev | Monomethyl auristatin E (MMAE) | Cleavable | Approved |
| Tisotumab vedotin | Tivdak | Monomethyl auristatin E (MMAE) | Cleavable | Approved |
| Moxetumomab pasudotox | Lumoxiti | Pseudomonas exotoxin (PE38) | Non-Cleavable | Approved |
| Loncastuximab tesirine | Zynlonta | SG3199 (pyrrolobenzodiazepine dimer) | Cleavable | Approved |
| Sacituzumab govitecan | Trodelyv | SN-38 (active metabolite of irinotecan) | Cleavable | Approved |
| Patritumab deruxtecan | N/A | Dxd (exatecan derivative) | Cleavable | In Clinical Trials |
| Telisotuzumab vedotin | Teliso-V | Monomethyl auristatin E (MMAE) | Cleavable | In Clinical Trials |
| Disitamab vedotin | RC48 | Monomethyl auristatin E (MMAE) | Cleavable | In Clinical Trials |
| Depatuxizumab mafodotin | Depatux-M | Monomethyl auristatin F (MMAF) | Non-Cleavable | In Clinical Trials |
| Vadastuximab talirine | N/A | SGD-1882 (pyrrolobenzodiazepine dimer) | Cleavable | In Clinical Trials |
| Camidanlumab tesirine | Cami-T | SG3199 (pyrrolobenzodiazepine dimer) | Cleavable | In Clinical Trials |
| Belantamab mafodotin | Blenrep | Monomethyl auristatin F (MMAF) | Non-Cleavable | Withdrawn |

Despite some responses of cancers to ADCs, resistance to treatment is observed in the clinic. Clinically observed mechanisms of resistance include downregulation or mutation of the ADC antibody target, failure of lysosomes to release the payload, activation of alternative survival pathways, and the overexpression of efflux transporters^[12, 13]^. Patients whose leukemic blasts had lower levels of CD33 were found to respond more poorly to treatment with gemtuzumab ozogamicin versus patients with blasts expressing higher levels^[14]^. Similarly, patients whose tumors were found to express a truncated form of HER2 that does not bind to trastuzumab, p95HER2, were found to be largely resistant to trastuzumab emtansine^[15]^. Decreased payload release in patient-derived samples has been linked to drug accumulation in the lysosomes due to increased lysosomal pH^[16]^. Finally, overexpression of the *ABCB1* gene which encodes the ATP-binding cassette (ABC) transporter P-glycoprotein (P-gp) has been inversely correlated with response to gemtuzumab ozogamicin in patients with AML^[14, 17]^. While *ad hoc* reports of ABC-mediated resistance of individual ADCs exist^[18, 19]^, no systematic examination of the relationship between toxin sensitivity and ABC transporter expression has been reported.

The objective of this study was to determine whether several commonly used ADC payloads were substrates for transport by ABC multidrug transporters, with the goal of identifying cytoxic agents that were not subject to transport out of the cell. P-gp substrates were identified by comparing compound activity between paired sets of low-expressing (sensitive) and over-expressing (resistant) cell lines for both P-gp and ABCG2. Four paired cell sets were utilized – a drug-selected and transfected set for each transporter. Hits were confirmed in cells transfected to express P-gp, ABCG2, or MRP1 (encoded by *ABCC1*). These results should aid investigators in designing ADCs that are not subject to ABC transporter-mediated resistance.

## Methods

### Cell Lines

HEK-293 cells were transfected with empty vector (pcDNA) or vector containing the human *ABCB1* (MDR-19), *ABCG2* (R-5), or *ABCC1* gene (MRP1) and were maintained in EMEM with 2 mg/mL G418 added to maintain transporter expression^[20]^. The HeLa derivative KB-3-1 and the P-gp overexpressing KB-8-5-11 subline were maintained in DMEM; KB-8-5-11 additionally received 100 ng/mL colchicine^[21]^. OVCAR8, NCI-ADR-RES, and H460 parental cells were obtained from the Division of Cancer Treatment and Diagnosis Tumor Repository, National Cancer Institute at Frederick, MD and were cultured in RPMI. The ABCG2 expressing subline H460 MX20 was generated by selecting H460 cells in 20 nM mitoxantrone^[22]^. Media in all cases was supplemented with 10% FBS and 1% penicillin-streptomycin. We also performed screening against a panel of 99 cancer cell lines as outlined in Table S1. Cell line source and culture conditions are reported in Table S1; medium in all cases contained 1% penicillin-streptomycin.

### High-throughput screen

We explored 27 common ADC payloads, listed in Table 2, in two orthogonal, paired cell line sets—a drug selected and transfected set for each transporter—to identify substrates of the ABC transporters P-gp and ABCG2. As noted in Table 2, compounds were sourced from Levena Biopharma (San Diego, CA), Selleck Chem (Houston, TX), ChemieTek (Indianapolis, IN), or Microsource Discovery Systems (Gaylordsville, CT). Screening was performed as previously described^[23]^. Briefly, cells were plated at a density of 500 cells/well in 1536 well plates with compounds added using a 1536-head pin tool (Kalypsys, San Diego CA). Cells were incubated with cytotoxic drugs and processed with CellTiter Glo reagent (Promega, Madison, WI) to determine viability after 48 hours. P-gp substrates were identified by comparing compound activity between paired cell line sets of low (sensitive) and high (resistant) transporter expression.

**Table 2:** ADC payloads tested in the high throughput screen.

| Compound | Vendor | Mechanism of action |
| --- | --- | --- |
| N-Acetyl-Calicheamicin gamma1 | Levena Biopharma | DNA damage |
| DM1 | Levena Biopharma | Microtubule disruptor |
| PBD Dimer (SGD-1882) | Levena Biopharma | DNA damage |
| Monomethyl Auristatin E (MMAE) | Levena Biopharma | Microtubule disruptor |
| SN-38 | ChemieTek | Topoisomerase inhibition |
| DM4 | Levena Biopharma | Microtubule disruptor |
| alpha-Amanitin | Levena Biopharma | RNA polymerase inhibitor |
| Duocarmycin DM | Levena Biopharma | DNA alkylation |
| Monomethyl Auristatin F (MMAF) | Levena Biopharma | Microtubule disruptor |
| PNU 159682 | Levena Biopharma | Topoisomerase inhibition |
| Maytansinol | Levena Biopharma | Microtubule disruptor |
| Duocarmycin SA | Levena Biopharma | DNA alkylation |
| Monomethyl Dolastatin 10 | Levena Biopharma | Microtubule disruptor |
| Vinblastine Sulfate | Microsource Discovery | Microtubule disruptor |
| Ansamitocin P3 | Levena Biopharma | Microtubule disruptor |
| Calicheamicin gamma1 | Levena Biopharma | DNA damage |
| Nemorubicin | Levena Biopharma | Topoisomerase inhibition |
| Doxorubicin | Selleck Chem | Topoisomerase inhibition |
| Tubulysin IM-2 | Levena Biopharma | Microtubule disruptor |
| Auristatin F | Levena Biopharma | Microtubule disruptor |
| Camptothecin | Selleck Chem | Topoisomerase inhibition |
| Dasatinib | Selleck Chem | Tyrosine kinase inhibitor |
| Duocarmycin MA | Levena Biopharma | DNA alkylation |
| Duocarmycin TA | Levena Biopharma | DNA alkylation |
| Tubulysin M | Levena Biopharma | Microtubule disruptor |
| Auristatin E | Levena Biopharma | Microtubule disruptor |
| Dolastatin 10 | Levena Biopharma | Microtubule disruptor |

### Synergy experiments

Known inhibitors of the P-gp and ABCG2 pumps, tariquidar (1 µM) and Ko143 (5 µM) respectively, were co-administered with the payloads. Inhibition of efflux can restore intracellular accumulation; thus, re-sensitization of resistant cells confirms cytotoxic compounds are transporter substrates. Cells were incubated with drugs at various concentrations and processed with CellTiter Glo reagent (Promega, Madison, WI) to determine viability after 48 hours.

### Data analysis

To evaluate compound activity in high-throughput screening (HTS) and synergy experiments, concentration-response curves (CRCs) were generated for each sample by plotting normalized response data against compound concentration. These curves were modeled using a four-parameter logistic regression, which provided key pharmacological parameters such as the half-maximal inhibitory concentration (IC₅₀) and maximal response (efficacy)^[24]^. While many compounds showed well-defined, sigmoidal dose– response behavior with both upper and lower asymptotes, others displayed atypical or poor-quality CRCs, including shallow slopes, incomplete asymptotes, or responses derived from only a single active concentration point. Such cases were classified as low-confidence actives due to limited or ambiguous activity profiles. Furthermore, a compound’s area-under-the-curve (AUC) calculated based on the screening data analysis and curve fittings served as an integrated measure of compound activity, capturing both potency and efficacy, and was utilized for direct comparison of activity outcome across different cell lines and experimental conditions^[23]^.

### Confirmatory experiments with screen hits

Three-day cytotoxicity assays were performed using various payloads including other members of the PBD dimer class, as well as the ADC mirvetuximab soravtansine (obtained from MedChemExpress, Monmouth Junction, NJ), with pcDNA (empty vector transfected), MDR-19 (*ABCB1* transfected), R-5 (*ABCG2* transfected) and MRP1 (*ABCC1* transfected) cells. Briefly, cells were plated (5000 cells/well) in 96-well plates and allowed to attach overnight after which the desired payloads or the ADC at various concentrations were added and incubated with cells for three days. CellTiter Glo reagent (Promega) was then used according to the manufacturer’s directions to assess GI_50_ values.

## Results

### A High throughput screen (HTS) identifies ADC payloads as substrates of P-gp and ABCG2

We evaluated 27 common ADC payloads in two orthogonal, paired cell sets to identify substrates of the ABC transporters P-gp and ABCG2 (Table 2). Cytotoxic payloads were tested against two pairs of cell lines, one parental and one transporter-expressing line. For P-gp, the pairs were: (1) the parental KB 3-1 human adenocarcinoma cell line and its colchicine-selected, P-gp-overexpressing sub-line KB 8-5-11; and (2) the HEK pcDNA human embryonic kidney cell line (transfected with an empty vector plasmid control) and its *ABCB1* stably transfected sub-line MDR-19. For ABCG2 the pairs were: (1) the parental H460 human lung carcinoma cell line and its mitoxantrone-selected, ABCG2-overexpressing sub-line H460 MX20; and (2) the same HEK pcDNA parent cell line as a comparator for the *ABCG2*-transfected HEK cell line, R-5. Transporter-specific efflux was demonstrated by testing all toxins for sensitization of transporter-expressing cells in the presence of either the P-gp inhibitor tariquidar or the ABCG2 inhibitor Ko143. As such, six cell line conditions were used to evaluate toxins as P-gp substrates, and six cell line conditions were also used for evaluation of ABCG2 substrates.

Area under the dose response curve (AUC) values were calculated for each cell line with all payloads in the presence or absence of a specific inhibitor as previously described^[23]^ and are displayed in the heat maps in Figure 1. In the heatmaps, red denotes high cell death and thus a more potent compound, while blue denotes less cell death. In Figure 1A, cells expressing P-gp, when compared to parental cells, were found to be less sensitive (*i.e.,* resistant) to several ADC payloads, including calicheamicin gamma1, dolostatin 10, MMAE, tubulysin M, DM1, and DM4. When the P-gp positive lines were incubated with the inhibitor tariquidar, resistance to calicheamicin gamma1, dolostatin 10, MMAE, tubulysin M, DM1 and DM4 was reversed. In the heat map, unsupervised clustering showed that the KB 8-5-11 line treated with tariquidar clustered with the parent KB-3-1 line and the MDR-19 line with tariquidar clustered with pcDNA transfected cells, while the P-gp expressing cells cluster separately. The cytotoxicity of nemorubicin, its metabolite PNU-159682, and the duocarmycin compounds appeared to be unaffected by P-gp levels.

**Figure 1:**
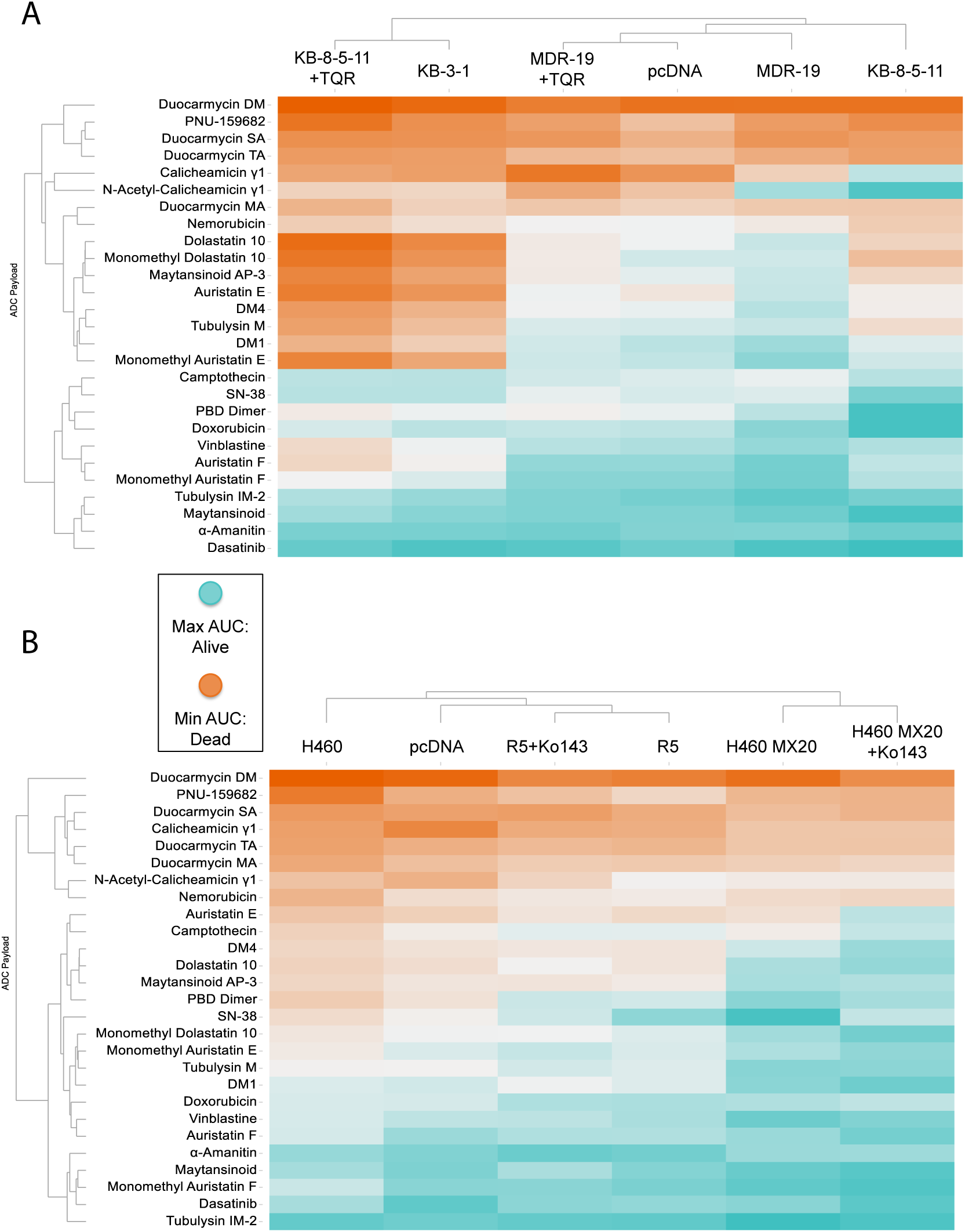
Heatmap of ADC payload activity across multidrug-resistant and control cell lines. **(A)** Unsupervised clustering of area under the curve (AUC) values from dose-response viability assays of a panel of cytotoxic agents tested against parental, multidrug-resistant, and genetically modified KB and MDR-19 cell lines. Cell lines include KB-3-1 (parental), KB-8-5-11 (ABCB1 overexpressing), MDR-19 (ABCB1-overexpressing), and their vector controls (pcDNA) or P-gp–inhibited counterparts (+TQR, tariquidar). **(B)** Unsupervised clustering of AUC values from the same compound panel tested in H460 (non-small cell lung cancer) parental cells, MDR sublines (H460 MX20 and H460 MX20+Ko143), and control or ABCG2-overexpressing derivatives (R5, R5+Ko143). Color scale reflects compound potency: orange indicates stronger cell killing (more cell death; greater sensitivity), while cyan indicates weaker cell killing (less cell death; more resistance). Compounds are grouped by activity outcome as indicated on the left dendrogram. Clustering was performed based on Euclidean distances of AUC profiles across cell lines.

As shown in in Figure 1B, few compounds appeared to be substrates of ABCG2, with the known substrate SN-38 being the most prominent compound that was subject to transport. Addition of the ABCG2 inhibitor Ko143 did not appear to sensitize cells, as R-5 cells with Ko143 clustered with R-5 cells, and H460 MX20 cells with Ko143 clustered with H460 MX20 cells, rather than with the corresponding parental lines. In line with efflux by ABCG2, the addition of Ko143 reversed resistance to SN-38.

We also calculated the differences in AUC (deltaAUC) values between the parental and resistant lines and between the resistant lines in the absence or presence of inhibitor, as previously described^[23]^. Comparing deltaAUC values between the pcDNA/MDR-19 and MDR-19+tariquidar pairs and the KB 3-1/KB 8-5-11 and KB 8-5-11+tariquidar pairs, we found good correlation, with r^2^ values of 0.7207 and 0.9175, respectively (Supplementary Figure 1). The results with P-gp contrasted with those of ABCG2, as few compounds were identified as ABCG2 substrates; the deltaAUC values for the pcDNA/R-5 and R-5 Ko143/R-5 pairs as well as the H460/H460MX20 and H460 Ko143/H460MX20 pairs showed poor correlation, with r^2^ values of 0.32 and 0.31 respectively (Supplementary Fig 1). Thus, more payloads appeared to be transported by P-gp than ABCG2.

### Validation of HTS hits

We validated selected screening results, adding a cell line (MRP1) developed by transfecting HEK293 cells with a plasmid containing the *ABCC1* gene. (Figure 2, Table 3). We included more camptothecin derivatives that serve as ADC payloads. Noting that PDB dimers were substrates of P-gp in the initial screen, we also added more PDB dimer payloads in our follow follow-up. Consistent with data from the screen, MMAE, DM4, calicheamicin gamma1, and DM1 were confirmed as avid substrates of P-gp and we noted a small degree of resistance conferred to these payloads by ABCG2 (Figure 2). MMAF was not a substrate of any of the transporters, although it was less potent than the related compound MMAE (Table 3). We also confirmed that nemorubicin and PNU159682 were not substrates of any of the transporters examined, making these ideal ADC payloads; PNU159682 is exquisitely toxic, with GI_50_ values in the picomolar range (Table 3).

**Figure 2:**
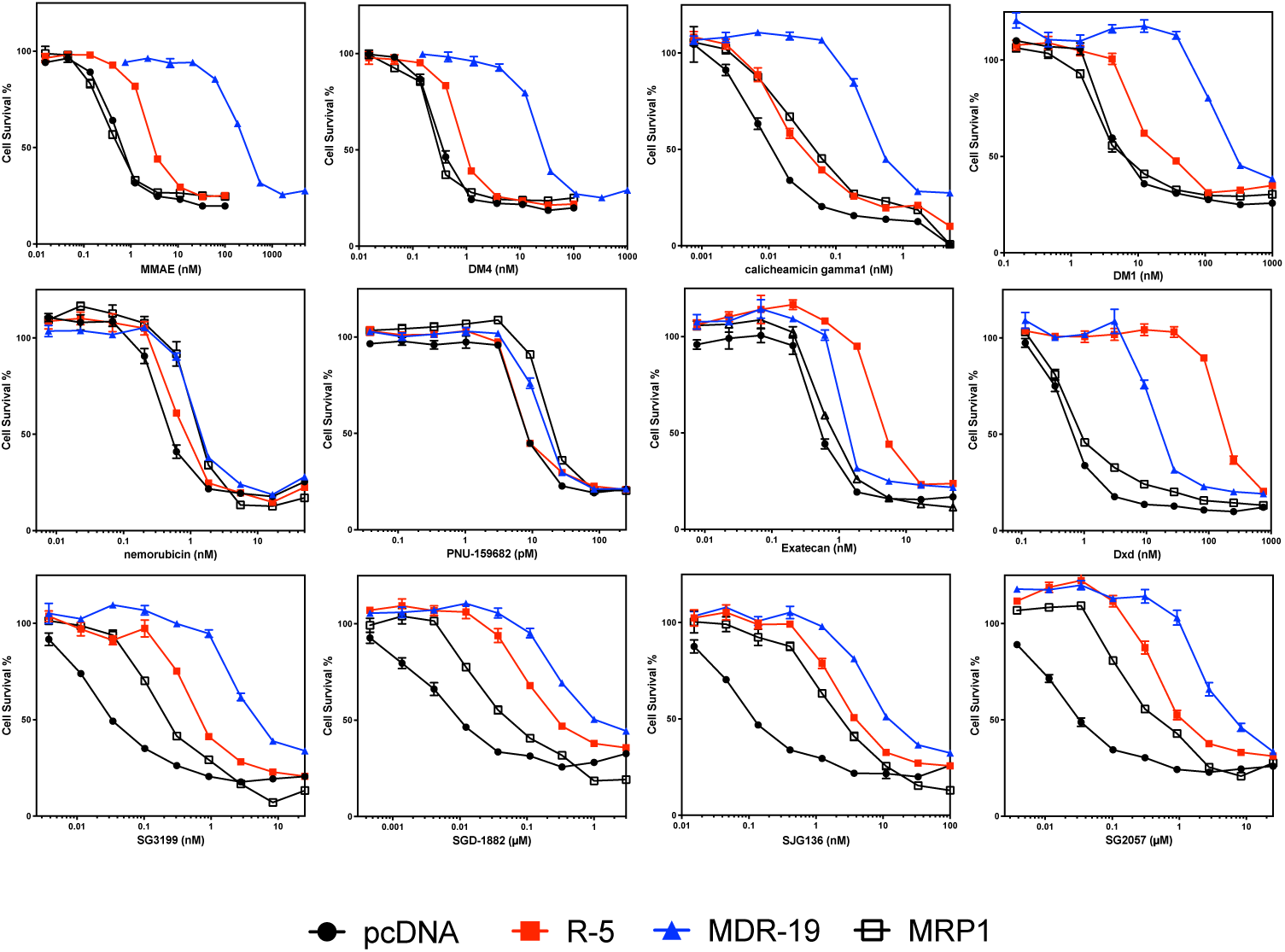
Confirmatory cytotoxicity assays with screen hits additional ADC payloads. Three-day cytotoxicity assays were performed with the noted compounds as described in the Materials and Methods using HEK-293 cells transfected with empty vector (pcDNA, black dot), or vectors containing *ABCG2* (R5, red square), *ABCB1* (MDR-19, blue triangle), or *ABCC1* (MRP1, black box). Results from one of three independent experiments are shown. Cytotoxicity data are summarized in Table 3.

**Table 3:** Cross-resistance profile to select ADC payloads in cells expressing P-gp, ABCG2 or MRP1.

| Drug | pcDNA | MDR19 | RR | R5 | RR | MRP1 | RR |
| --- | --- | --- | --- | --- | --- | --- | --- |
| DM1 (nM) | 7.8±1.4 | 410±120 | 52 | 35±4.1 | 4.5 | 9.5±3.7 | 1.2 |
| DM4 (nM) | 0.43±0.075 | 42±20 | 96 | 1.4±0.49 | 3.3 | 0.58±0.37 | 1.3 |
| MMAE (nM) | 0.90±0.23 | 360±110 | 400 | 5.1±1.9 | 5.7 | 1.2±0.80 | 1.3 |
| MMAF (μM) | 0.99±.097 | 2.0±1.7 | 2.0 | 0.99±.097 | 1.0 | 1.0±0.39 | 1.0 |
| Calicheamicin gamma1 | 0.010±0.0042 | 0.73±0.41 | 71 | 0.029±0.0047 | 2.9 | 0.028±0.017 | 2.7 |
| Exatecan (nM) | 0.59±0.19 | 1.4±0.15 | 2.4 | 5.0±0.80 | 8.5 | 1.0±0.19 | 1.8 |
| Deruxtecan (nM) | 0.82±0.18 | 17±1.2 | 20 | 190±29 | 240 | 1.1±0.40 | 1.4 |
| Nemorubicin (nM) | 0.76±0.31 | 1.5±0.15 | 2.0 | 1.1±0.27 | 1.4 | 1.6±0.21 | 2.2 |
| PNU-159682 (pM) | 13±9.6 | 16±3.7 | 1.2 | 13±8.4 | 1.0 | 19±8.6 | 1.4 |
| SJG136 (nM) | 0.098±0.019 | 16±6.2 | 170 | 4.6±0.68 | 47 | 1.2±1.1 | 12 |
| SGD-1882 (nM) | 0.0066±0.0028 | 0.85±0.34 | 130 | 0.22±0.093 | 33 | 0.095±0.036 | 14 |
| SG2057 (nM) | 0.025±0.0087 | 5.7±1.2 | 230 | 0.73±0.34 | 29 | 0.27±0.20 | 11 |
| SG3199 (nM) | 0.045±0.011 | 5.6±1.6 | 120 | 0.90±0.31 | 20 | 0.21±0.026 | 4.6 |

Results are mean GI50 values +/- standard deviation. Relative resistance (RR) value was computed by dividing the GI50 value for the lines expressing the ABC transporters by the empty vector (pcDNA) line. Results are from 3 independent experiments.

Overexpression of ABCG2 conferred about 10-fold resistance to the camptothecin derivative exatecan, much less than the 96-fold resistance for SN-38 we previously reported for ABCG2 transfected cells^[25]^. This is consistent with previous reports demonstrating that ABCG2 confers less resistance to exatecan compared to other camptothecin derivatives that are ABCG2 substrates^[26]^. However, the exatecan derivative Dxd (deruxtecan) was readily transported by ABCG2 and was also a substrate for P-gp (Figure 2, Table 3).

As our screen originally contained only one member of the PBD dimer family (SGD-1882), we expanded the number of PDB dimer payloads examined. We found that HEK293 cells expressing any of the three transporters conferred some resistance to all PDB dimers examined: SG3199, SGD-1882, SJG136 (also known as SG2000), and SG2057. P-gp overexpression was found to confer the highest levels of resistance, followed by ABCG2 and finally MRP1 (Figure 3, Table 3).

**Figure 3:**
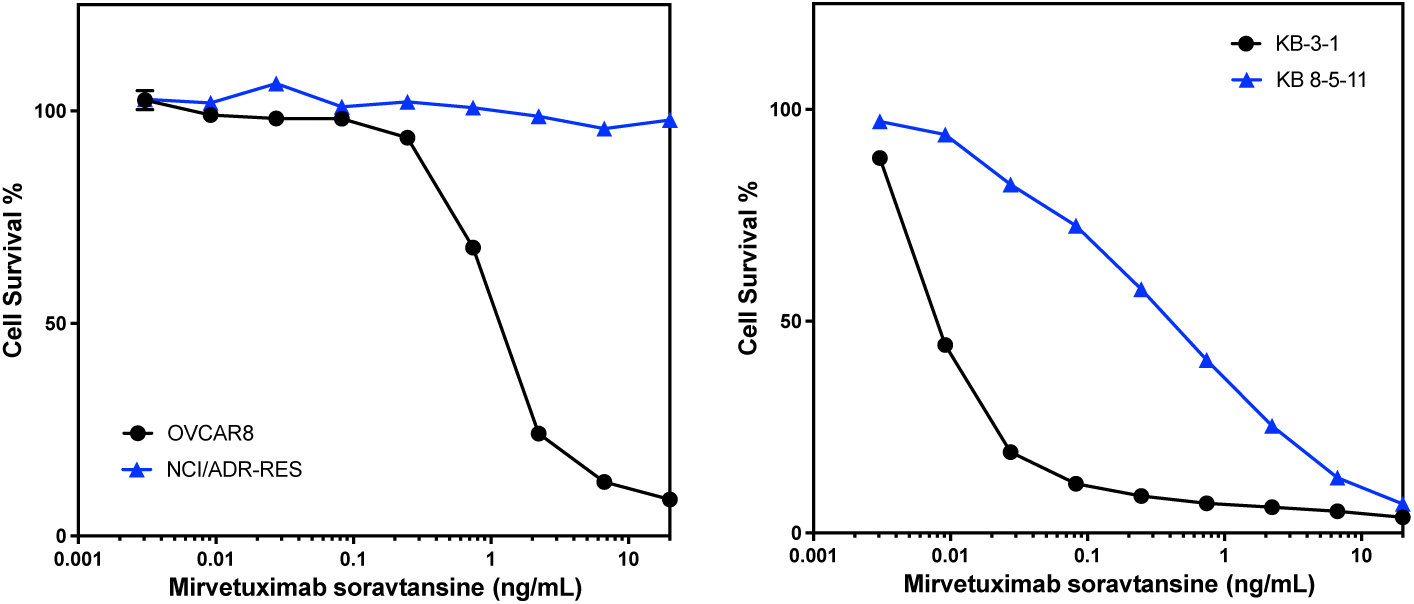
P-gp overexpression confers resistance to treatment with mirvetuximab soravtansine. Three-day cytotoxicity assays were performed with mirvetuximab soravtansine as described in the Materials and Methods using the OVCAR8/NCI/ADR-RES pair (left graph) or the KB-3-1/KB 8-5-11 pair (right graph). Results from one of three independent experiments are shown.

### P-gp overexpression confers resistance to mirvetuximab soravtansine

The screening approach described here evaluated small molecule cytotoxic drugs employed as payloads in experimental and therapeutic ADCs. Examining how drug transporter-mediated susceptibility confers resistance to an ADC (where the toxin is liberated from the antibody within the cell and subsequently effluxed) is important for extrapolating our results. We thus tested the effect of P-gp expression on the sensitivity of cultured cells to an ADC. Mirvetuximab soravtansine (MIRV) is an ADC that was approved by the FDA in 2022 for the treatment of folate receptor alpha-positive, platinum-resistant ovarian, fallopian tube or peritoneal cancer^[10]^. The toxic payload is DM4, which we found to be a P-gp substrate in our screen. Testing MIRV with the KB3-1/KB-8-5-11 pair used in our screening assay and the OVCAR8/NCI-ADR-RES cell line pair, we found that the P-gp overexpressing lines KB 8-5-11 and NCI-ADR-RES were resistant to treatment with MIRV compared to the parental lines. These results confirm that screening of isolated payloads for susceptibility to P-gp efflux translates to cellular resistance to an ADC carrying a P-pg-substrate toxin. These data suggest that P-gp may contribute to resistance to MIRV in the clinic.

### Duocarmycins and PNU-159682 are highly toxic to a broad range of cancer cell lines

We next explored the toxicity of the various payloads across a broad range of cell lines, including lines from cancers originating from the skin, kidney, pancreas, lung and blood. We excluded five payloads from our original list of 27—vinblastine, SN-38, camptothecin, dasatinib and doxorubicin—as these compounds have been well characterized. As shown in Figure 4, hematologic cancers were sensitive to nearly all of the compounds tested, while cancers of the lung and pancreas tended to be less sensitive. The duocarmycins and PNU-159682 were uniformly toxic across the cell line panel. Given that the duocarmycins and PNU-159682 were not found to be substrates of ABC transporters, these compounds should be utilized as ADC payloads in the future.

**Figure 4:**
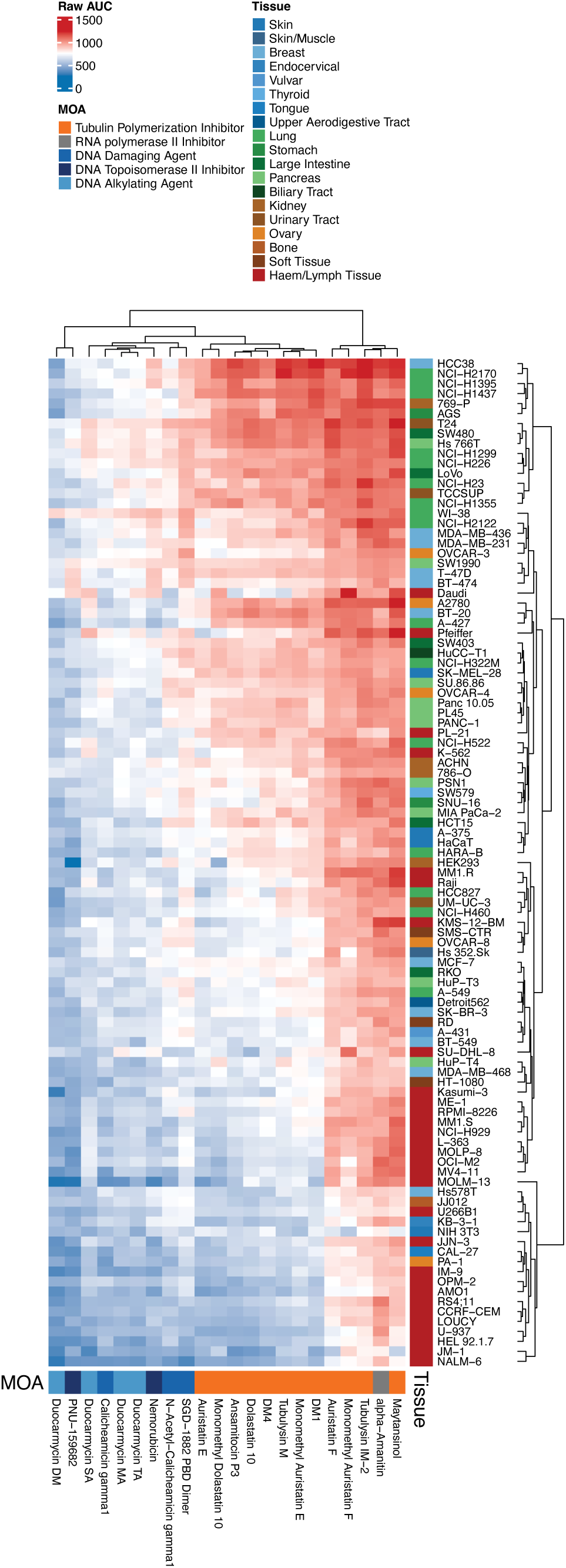
Differential cytotoxicity of ADC payloads across cancer cell lines from diverse tissue origins. Heatmap showing AUC values from dose–response viability assays for a panel of ADC payload compounds tested across a large set of human cancer cell lines. Each row represents a unique cancer cell line, annotated by tissue of origin (colored bar on the right-hand side), and each column corresponds to an ADC payload compound, annotated by mechanism of action (MOA) on the bottom of the heatmap. AUC values are color-coded, with red indicating higher AUC (lower sensitivity/more resistant) and blue indicating lower AUC (higher sensitivity/more cytotoxic effect). Clustering (both rows and columns) was performed using hierarchical methods based on compound response profiles.

## Discussion

We systematically screened 27 commonly used ADC payloads (cytotoxic drugs) in two orthogonal, paired cell sets to identify substrates of the ABC transporters P-gp and ABCG2. Of the studied DNA damaging agents, calicheamicin analogs, a PDB dimer, SN-38, and doxorubicin were identified as substrates of the ABC transporter P-gp. Of the tubulin-targeting agents, vinblastine and all auristatin and maytansinoid derivatives were strong P-gp substrates, whereas MMAF was somewhat less susceptible to transport. Notably, the following compounds were found to not be P-gp substrates: duocarmycin analogs, camptothecin, nemorubicin, PNU-159682, and α-amanitin. SN-38 was identified as a substrate of ABCG2, but no other cytotoxic payloads were observed to be transported by ABCG2. Some cytotoxic drugs exhibited decreased sensitivity to the drug-selected H460-MX20 cell line, but this sensitivity was not reversed by co-dosing with the known ABCG2 inhibitor Ko143, suggesting that the difference was not due to ABCG2 overexpression.

*In vitro* studies and *in vivo* mouse models have demonstrated that overexpression of ABC transporters can cause resistance to some ADCs. Early studies with gemtuzumab ozogamicin demonstrated that P-gp and possibly MRP1 (ABCC1) played a role in resistance. Naito *et al.* found that leukemia cell lines that overexpressed P-gp were resistant to treatment with gemtuzumab ozogamicin and that combining the ADC with P-gp inhibitors such as valspodar or biricodar reversed resistance^[27]^. In the MRP1-overexpressing cell line HL-60/ADR, addition of the MRP1 inhibitor MK-571 was found to increase sensitivity to gemtuzumab ozogamicin, suggesting that MRP1 was yet another resistance mechanism to the ADC^[28]^; however, overexpression of ABCG2 does not appear to cause resistance^[29]^. Studies using cell lines expressing P-gp, MRP1 or ABCG2 with the CD33-targeting ADC AVE9633, which has the maytansine DM4 as its toxic payload, showed that only P-gp could confer resistance^[18]^. Repeated treatment with the ADC N41mab-vcMMAE, which targets nectin-4 and has MMAE as its payload, in mice xenografted with the SUM190 breast cancer cell line induced refractory tumors which were found to overexpress *ABCB1* as the mechanism of resistance^[30]^. An ADC targeting the delta-like non-canonical Notch ligand 1 protein, ADCT-701, which has the PDB dimer SG3199 as the payload, was found to be less effective in adrenocortical cancer cell lines and organoids with high expression of P-gp, and the free drug was also reported to be a substrate of P-gp^[31]^. The findings of these studies agree with our results confirming that calicheamicin, DM4, MMAE and SG3199 are substrates of P-gp.

Overexpression of P-glycoprotein has also emerged as a marker of resistance in a subset of patients who have been treated with gemtuzumab ozogamicin and brentuximab vedotin. Early studies examining leukemic blasts from patients with resistant disease demonstrated that P-gp and MRP1 expression could lead to resistance^[17, 28, 32]^. P-gp expression was found to inversely correlate with clinical response to gemtuzumab ozogamicin in patients with AML^[14]^. Similarly, a study examining a small cohort of lymphoma patients resistant to gemtuzumab vedotin reported P-gp overexpression in a patient with Hodgkin Lymphoma^[19]^. A case report of a patient with bladder cancer whose disease had progressed after treatment with enfortumab vedotin, an ADC which targets nectin-4, reported high levels of P-gp in the resistant tumor^[33]^.

These initial studies suggested that the addition of a P-gp inhibitor might be beneficial to patients whose cancer expresses P-gp, leading to some clinical trials combining P-gp inhibitors with ADC treatment. The P-gp inhibitor zosuquidar was found to reverse resistance to gemtuzumab ozogamycin in *ex vivo* studies using P-gp positive blasts obtained from patients with resistant disease^[34]^. In a clinical trial combining zosuquidar with gemtuzumab ozogamicin, greater overall survival was noted in patients with P-gp-positive resistant disease^[35]^. A small clinical study combining cyclosporine with gemtuzumab ozogamicin in patients with resistant disease led to an increased overall and complete response rate^[36]^. However, the final findings from a larger trial were less positive, mostly due to toxicity from the use of cyclosporine A as the P-gp inhibitor^[37]^. The disappointing results from this later trial highlight the importance of P-gp inhibitor choice when designing combination trials.

## Conclusion

We show that the payloads of some FDA approved ADCs are strong substrates for P-gp, suggesting that active transport and efflux of a released payload may play a role in acquired resistance to clinical ADCs. These effects may also increase exposure of the cytotoxic payload to systemic tissues and could contribute to *in vivo* off-target toxicities. We identified ADC payloads that were not substrates of either P-gp or ABCG2—notably the duocarmycin series and PNU-159682, which were among the most cytotoxic of all the toxins studied in a panel of 99 cancer cell lines—and suggest that these compounds be prioritized as future ADC payloads due to the potential for reduced susceptibility to transporter-mediated acquired resistance.

## Author contributions

Jacob S. Roth: data curation, formal analysis, investigation, visualization, writing— original draft preparation, writing—review and editing

Hui Guo: formal analysis, investigation, visualization

Lu Chen: formal analysis, investigation, visualization

Min Shen: formal analysis, investigation, visualization, writing—original draft preparation, writing—review and editing

Omotola Gbadagesin: data curation, formal analysis, investigation, writing—review and editing

Robert W. Robey: data curation, visualization, investigation, writing—original draft preparation, writing—review and editing

Michael M. Gottesman: supervision, writing—review and editing

Matthew D. Hall: conceptualization, resources, supervision, writing—original draft preparation, writing—review and editing

## Ethical approval and consent to participate

Not applicable.

## Consent for publication

Not applicable.

## References

1. He J, Zeng X, Wang C, Wang E and Li Y. Antibody-drug conjugates in cancer therapy: mechanisms and clinical studies. MedComm (2020) 2024;5:e671

2. Strebhardt K and Ullrich A. Paul Ehrlich’s magic bullet concept: 100 years of progress. Nat Rev Cancer 2008;8:473–80

3. Sievers EL, Larson RA, Stadtmauer EA, Estey E, Lowenberg B, Dombret H, Karanes C, Theobald M, Bennett JM, Sherman ML, Berger MS, Eten CB, Loken MR, van Dongen JJ, Bernstein ID, Appelbaum FR and Mylotarg Study G. Efficacy and safety of gemtuzumab ozogamicin in patients with CD33-positive acute myeloid leukemia in first relapse. J Clin Oncol 2001;19:3244–54

4. Senter PD and Sievers EL. The discovery and development of brentuximab vedotin for use in relapsed Hodgkin lymphoma and systemic anaplastic large cell lymphoma. Nat Biotechnol 2012;30:631–7

5. Amiri-Kordestani L, Blumenthal GM, Xu QC, Zhang L, Tang SW, Ha L, Weinberg WC, Chi B, Candau-Chacon R, Hughes P, Russell AM, Miksinski SP, Chen XH, McGuinn WD, Palmby T, Schrieber SJ, Liu Q, Wang J, Song P, Mehrotra N, Skarupa L, Clouse K, Al-Hakim A, Sridhara R, Ibrahim A, Justice R, Pazdur R and Cortazar P. FDA approval: ado-trastuzumab emtansine for the treatment of patients with HER2-positive metastatic breast cancer. Clin Cancer Res 2014;20:4436–41

6. Modi S, Jacot W, Yamashita T, Sohn J, Vidal M, Tokunaga E, Tsurutani J, Ueno NT, Prat A, Chae YS, Lee KS, Niikura N, Park YH, Xu B, Wang X, Gil-Gil M, Li W, Pierga JY, Im SA, Moore HCF, Rugo HS, Yerushalmi R, Zagouri F, Gombos A, Kim SB, Liu Q, Luo T, Saura C, Schmid P, Sun T, Gambhire D, Yung L, Wang Y, Singh J, Vitazka P, Meinhardt G, Harbeck N, Cameron DA and Investigators DE-BT. Trastuzumab Deruxtecan in Previously Treated HER2-Low Advanced Breast Cancer. N Engl J Med 2022;387:9–20

7. Meric-Bernstam F, Makker V, Oaknin A, Oh DY, Banerjee S, Gonzalez-Martin A, Jung KH, Lugowska I, Manso L, Manzano A, Melichar B, Siena S, Stroyakovskiy D, Fielding A, Ma Y, Puvvada S, Shire N and Lee JY. Efficacy and Safety of Trastuzumab Deruxtecan in Patients With HER2-Expressing Solid Tumors: Primary Results From the DESTINY-PanTumor02 Phase II Trial. J Clin Oncol 2024;42:47–58

8. Caimi PF, Ai W, Alderuccio JP, Ardeshna KM, Hamadani M, Hess B, Kahl BS, Radford J, Solh M, Stathis A, Zinzani PL, Havenith K, Feingold J, He S, Qin Y, Ungar D, Zhang X and Carlo-Stella C. Loncastuximab tesirine in relapsed or refractory diffuse large B-cell lymphoma (LOTIS-2): a multicentre, open-label, single-arm, phase 2 trial. Lancet Oncol 2021;22:790–800

9. Yu EY, Petrylak DP, O’Donnell PH, Lee JL, van der Heijden MS, Loriot Y, Stein MN, Necchi A, Kojima T, Harrison MR, Hoon Park S, Quinn DI, Heath EI, Rosenberg JE, Steinberg J, Liang SY, Trowbridge J, Campbell M, McGregor B and Balar AV. Enfortumab vedotin after PD-1 or PD-L1 inhibitors in cisplatin-ineligible patients with advanced urothelial carcinoma (EV-201): a multicentre, single-arm, phase 2 trial. Lancet Oncol 2021;22:872–82

10. Moore KN, Angelergues A, Konecny GE, Garcia Y, Banerjee S, Lorusso D, Lee JY, Moroney JW, Colombo N, Roszak A, Tromp J, Myers T, Lee JW, Beiner M, Cosgrove CM, Cibula D, Martin LP, Sabatier R, Buscema J, Estevez-Garcia P, Coffman L, Nicum S, Duska LR, Pignata S, Galvez F, Wang Y, Method M, Berkenblit A, Bello Roufai D, Van Gorp T, Gynecologic Oncology Group P and the European Network of Gynaecological Oncological Trial G. Mirvetuximab Soravtansine in FRalpha-Positive, Platinum-Resistant Ovarian Cancer. N Engl J Med 2023;389:2162–74

11. Xi M, Zhu J, Zhang F, Shen H, Chen J, Xiao Z, Huangfu Y, Wu C, Sun H and Xia G. Antibody-drug conjugates for targeted cancer therapy: Recent advances in potential payloads. Eur J Med Chem 2024;276:116709

12. Jiang M, Li Q and Xu B. Spotlight on ideal target antigens and resistance in antibody-drug conjugates: Strategies for competitive advancement. Drug Resist Updat 2024;75:101086

13. Loganzo F, Sung M and Gerber HP. Mechanisms of Resistance to Antibody-Drug Conjugates. Mol Cancer Ther 2016;15:2825–34

14. Walter RB, Gooley TA, van der Velden VH, Loken MR, van Dongen JJ, Flowers DA, Bernstein ID and Appelbaum FR. CD33 expression and P-glycoprotein-mediated drug efflux inversely correlate and predict clinical outcome in patients with acute myeloid leukemia treated with gemtuzumab ozogamicin monotherapy. Blood 2007;109:4168–70

15. Scaltriti M, Rojo F, Ocana A, Anido J, Guzman M, Cortes J, Di Cosimo S, Matias-Guiu X, Ramon y Cajal S, Arribas J and Baselga J. Expression of p95HER2, a truncated form of the HER2 receptor, and response to anti-HER2 therapies in breast cancer. J Natl Cancer Inst 2007;99:628–38

16. Rios-Luci C, Garcia-Alonso S, Diaz-Rodriguez E, Nadal-Serrano M, Arribas J, Ocana A and Pandiella A. Resistance to the Antibody-Drug Conjugate T-DM1 Is Based in a Reduction in Lysosomal Proteolytic Activity. Cancer Res 2017;77:4639–51

17. Linenberger ML, Hong T, Flowers D, Sievers EL, Gooley TA, Bennett JM, Berger MS, Leopold LH, Appelbaum FR and Bernstein ID. Multidrug-resistance phenotype and clinical responses to gemtuzumab ozogamicin. Blood 2001;98:988–94

18. Tang R, Cohen S, Perrot JY, Faussat AM, Zuany-Amorim C, Marjanovic Z, Morjani H, Fava F, Corre E, Legrand O and Marie JP. P-gp activity is a critical resistance factor against AVE9633 and DM4 cytotoxicity in leukaemia cell lines, but not a major mechanism of chemoresistance in cells from acute myeloid leukaemia patients. BMC Cancer 2009;9:199

19. Chen R, Hou J, Newman E, Kim Y, Donohue C, Liu X, Thomas SH, Forman SJ and Kane SE. CD30 Downregulation, MMAE Resistance, and MDR1 Upregulation Are All Associated with Resistance to Brentuximab Vedotin. Mol Cancer Ther 2015;14:1376-84

20. Robey RW, Lin B, Qiu J, Chan LL and Bates SE. Rapid detection of ABC transporter interaction: potential utility in pharmacology. J Pharmacol Toxicol Methods 2011;63:217–22

21. Shen DW, Fojo A, Chin JE, Roninson IB, Richert N, Pastan I and Gottesman MM. Human multidrug-resistant cell lines: increased mdr-1 expression can precede gene amplification. Science 1986;232:643–45

22. Robey RW, Honjo Y, van de Laar A, Miyake K, Regis JT, Litman T and Bates SE. A functional assay for detection of the mitoxantrone resistance protein, MXR (ABCG2). Biochim Biophys Acta 2001;1512:171-82

23. Lee TD, Lee OW, Brimacombe KR, Chen L, Guha R, Lusvarghi S, Tebase BG, Klumpp-Thomas C, Robey RW, Ambudkar SV, Shen M, Gottesman MM and Hall MD. A High-Throughput Screen of a Library of Therapeutics Identifies Cytotoxic Substrates of P-glycoprotein. Mol Pharmacol 2019;

24. Inglese J, Auld DS, Jadhav A, Johnson RL, Simeonov A, Yasgar A, Zheng W and Austin CP. Quantitative high-throughput screening: a titration-based approach that efficiently identifies biological activities in large chemical libraries. Proc Natl Acad Sci U S A 2006;103:11473–8

25. Robey RW, Honjo Y, Morisaki K, Nadjem TA, Runge S, Risbood M, Poruchynsky MS and Bates SE. Mutations at amino acid 482 in the ABCG2 gene affect substrate and antagonist specificity. Br J Cancer 2003;89:1971–8

26. Ishii M, Iwahana M, Mitsui I, Minami M, Imagawa S, Tohgo A and Ejima A. Growth inhibitory effect of a new camptothecin analog, DX-8951f, on various drug-resistant sublines including BCRP-mediated camptothecin derivative-resistant variants derived from the human lung cancer cell line PC-6. Anticancer Drugs 2000;11:353-62

27. Naito K, Takeshita A, Shigeno K, Nakamura S, Fujisawa S, Shinjo K, Yoshida H, Ohnishi K, Mori M, Terakawa S and Ohno R. Calicheamicin-conjugated humanized anti-CD33 monoclonal antibody (gemtuzumab zogamicin, CMA-676) shows cytocidal effect on CD33-positive leukemia cell lines, but is inactive on P-glycoprotein-expressing sublines. Leukemia 2000;14:1436-43

28. Walter RB, Raden BW, Hong TC, Flowers DA, Bernstein ID and Linenberger ML. Multidrug resistance protein attenuates gemtuzumab ozogamicin-induced cytotoxicity in acute myeloid leukemia cells. Blood 2003;102:1466–73

29. Walter RB, Raden BW, Thompson J, Flowers DA, Kiem HP, Bernstein ID and Linenberger ML. Breast cancer resistance protein (BCRP/ABCG2) does not confer resistance to gemtuzumab ozogamicin and calicheamicin-gamma1 in acute myeloid leukemia cells. Leukemia 2004;18:1914–7

30. Cabaud O, Berger L, Crompot E, Adelaide J, Finetti P, Garnier S, Guille A, Carbuccia N, Farina A, Agavnian E, Chaffanet M, Goncalves A, Charafe-Jauffret E, Mamessier E, Birnbaum D, Bertucci F and Lopez M. Overcoming Resistance to Anti-Nectin-4 Antibody-Drug Conjugate. Mol Cancer Ther 2022;21:1227–35

31. Sun N-Y, Kumar S, Kim YS, Varghese D, Mendoza A, Nguyen R, Okada R, Reilly K, Widemann B, Pommier Y, Elloumi F, Dhall A, Patel M, Aber E, Contreras-Burrola C, Kaplan R, Martinez D, Pogoriler J, Hamilton AK, Diskin SJ, Maris JM, Robey RW, Gottesman MM, Del Rivero J and Roper N. Identification of DLK1, a Notch ligand, as an immunotherapeutic target and regulator of tumor cell plasticity and chemoresistance in adrenocortical carcinoma. bioRxiv 2024;2024.10.09.617077

32. Matsui H, Takeshita A, Naito K, Shinjo K, Shigeno K, Maekawa M, Yamakawa Y, Tanimoto M, Kobayashi M, Ohnishi K and Ohno R. Reduced effect of gemtuzumab ozogamicin (CMA-676) on P-glycoprotein and/or CD34-positive leukemia cells and its restoration by multidrug resistance modifiers. Leukemia 2002;16:813–9

33. Kotono M, Kijima T, Takada-Owada A, Okubo N, Kurashina R, Kokubun H, Uematsu T, Takei K, Ishida K and Kamai T. Increased expression of ATP-binding cassette transporters in enfortumab vedotin-resistant urothelial cancer. IJU Case Rep 2024;7:173–76

34. Tang R, Faussat AM, Perrot JY, Marjanovic Z, Cohen S, Storme T, Morjani H, Legrand O and Marie JP. Zosuquidar restores drug sensitivity in P-glycoprotein expressing acute myeloid leukemia (AML). BMC Cancer 2008;8:51

35. Marcelletti JF and Sikic BI. A clinical trial of zosuquidar plus gemtuzumab ozogamicin (GO) in relapsed or refractory acute myeloid leukemia (RR AML): evidence of efficacy based on leukemic blast P-glycoprotein functional phenotype. Cancer Chemother Pharmacol 2023;92:369–80

36. Chen R, Herrera AF, Hou J, Chen L, Wu J, Guo Y, Synold TW, Ngo VN, Puverel S, Mei M, Popplewell L, Yi S, Song JY, Tao S, Wu X, Chan WC, Forman SJ, Kwak LW, Rosen ST and Newman EM. Inhibition of MDR1 Overcomes Resistance to Brentuximab Vedotin in Hodgkin Lymphoma. Clin Cancer Res 2020;26:1034–44

37. Kambhampati S, Mei MG, Chen L, Puverel S, Chen R, Popplewell LL, Nikolaenko L, Peters L, Armenian S, Kwak LW, Rosen ST, Forman SJ and Herrera AF. Phase I Trial of Brentuximab Vedotin Plus Cyclosporine in Relapsed/Refractory Hodgkin Lymphoma. Clin Lymphoma Myeloma Leuk 2024;24:724–31 e1

