## Supplementary figures and images for "Identification of antibody-drug conjugate payloads which are substrates of ATP-binding cassette drug efflux transporters"

### Supplemental Figure 1

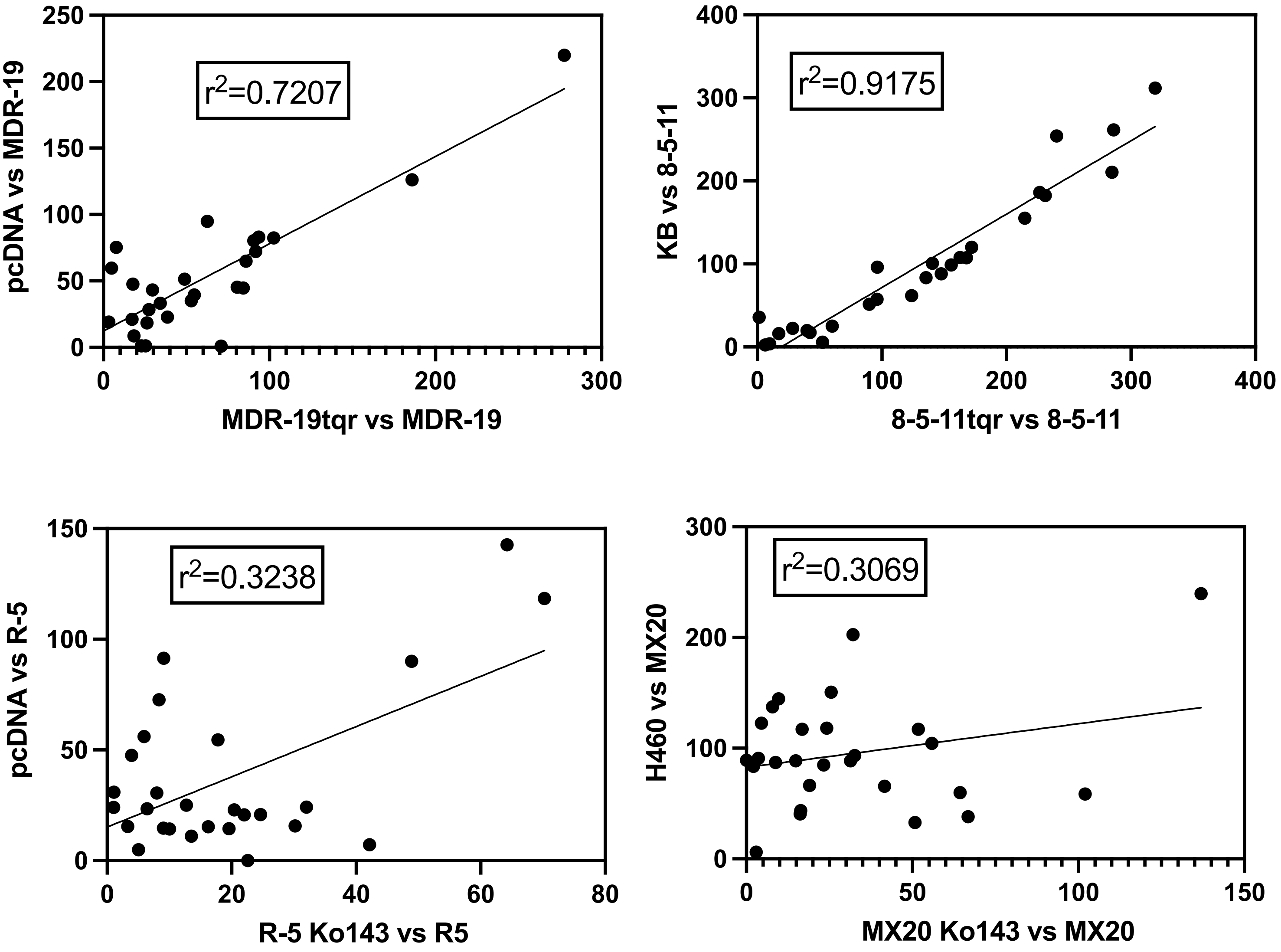
